# Altered gut microbiota and immunity defines *Plasmodium vivax* survival in *Anopheles stephensi*

**DOI:** 10.1101/774075

**Authors:** Punita Sharma, Jyoti Rani, Charu Chauhan, Seena Kumari, Sanjay Tevatiya, Tanwee Das De, Kailash C Pandey, Rajnikant Dixit

## Abstract

Blood feeding-enriched gut-microbiota boosts mosquitoes’ anti-*Plasmodium* immunity. Here, we ask how *Plasmodium vivax* alters microbiota, anti-*Plasmodial* immunity and impact tripartite *Plasmodium*-mosquito-microbiota interactions in the gut lumen. Using a metagenomics analysis, we predominantly detect *Elizabethkingia meningitis* and *Pseudomonas sps.* in naïve mosquitoes. Naïve blood fed gut shows a heightened presence of *Elizabethkingia*, *Pseudomonas* and *Serratia*. A parallel RNAseq analysis of blood-fed midguts identify *Elizabethkingia*-transcripts, which may have role in iron metabolism. Post, a *Plasmodium vivax* infected blood-meal, however, we do not detect bacterial until circa 36 hours. Intriguingly, transcriptional expression of a selected array of antimicrobial arsenal cecropins 1-2, defensin-1 and gambicin remains low during the first 36 hours–a time frame when ookinietes/early oocysts invade gut. We conclude during the preinvasive phase, *Plasmodium vivax* outcompetes midgut-microbiota. Suppression of important immune factors, likely due to altered microbiota, may enhance *Plasmodium vivax* survival. Additional finding of a novel *Wolbachia* association warrants further research to design ‘paratransgenesis’ tools for malaria control.

**Author Summary:** Successful malaria transmission relies on the competitive interactions of *Plasmodium* and mosquito’s tissue specific immune potential. Within 24hrs of blood meal gut-microbiota grows exponentially and lead to robust enhancement of mosquito immune response, which is detrimental to parasite survival and development. But the mechanism how *Plasmodium* manages to evade this pre-invasive immune barrier is not well known. We investigated the influence of tripartite gut-microbiome-parasite interaction on human malaria parasite *Plasmodium vivax* in its natural/native vector *Anopheles stephensi*. Surprisingly we found that infectious blood meal lead to dramatic suppression in gut-bacteria population, a plausible strategy of *P. vivax* ookinetes to avoid immune responses. Our study suggests that for its own survival *Plasmodium vivax* causes early suppression of bacterial population, possibly by scavenging Fe from the blood meal which is indispensable for bacterial growth. Disruption and manipulation of this gut-microbe-interaction may help to design new ‘paratransgenesis’ molecular tool for malaria control.

## Introduction

Blood meal is an essential requirement for the reproductive success of the adult female mosquitoes. Immediately after blood meal uptake, mosquitoes’ gut physiology undergoes complex modulation to facilitate rapid blood meal digestion and activation of vitellogenesis process (Attardo, Hansen et al. 2005; Richards, Anderson et al. 2012). The blood meal triggers proliferation of gut microbiome eliciting immune response (Gaio Ade, Gusmao et al. 2011; Romoli and Gendrin 2018) and once the blood meal digestion completed within first 30 hrs, the immune response apparently ceases to basal level (Pumpuni, Demaio et al. 1996; Das De, Sharma et al. 2018).

This gut immune response may indirectly affect the early development of *Plasmodium* when mosquitoes take infected blood (Dong, Manfredini et al. 2009; Smith, Vega-Rodriguez et al. 2014; Rodgers, Gendrin et al. 2017). Removal of gut microbes by antibiotic treatment enhances *Plasmodium* survival, but the exact mechanism that how *Plasmodium* manages its safe journey to gut and succeed to develop in the susceptible strains is yet to unravel (Noden, Vaughan et al. 2011). In the gut lumen an early tripartite interaction of gut-microbes-parasites may have a significant influence on the *Plasmodium* invasion dynamics (Simonetti 1996; Chavshin, Oshaghi et al. 2012; Chavshin, Oshaghi et al. 2014; Anglero-Rodriguez, Blumberg et al. 2016; Saraiva, Kang et al. 2016). But a great deal of understading that how parasite manages its survival during acute gut-microbe interaction is poorly understood (Romoli and Gendrin 2018). Once invaded the gut epithelial, the *Plasmodium* population undergoes several bottlenecks, reducing the oocysts load either to zero in naturally selected refractory mosquito strains, or a few oocysts in a susceptible mosquito vector species (Drexler, Vodovotz et al. 2008; Bennink, Kiesow et al. 2016).

Within 8-9 days post infection, the surviving oocysts rupture to millions of sporozoites, released in the hemolymph (Simonetti 1996). During free circulation, sporozoites compete to invade the salivary glands, and if not successful are rapidly cleared by the mosquito immune blood cells ‘hemocytes’ (Drexler, Vodovotz et al. 2008; Bennink, Kiesow et al. 2016; Belachew 2018; Das De, Sharma et al. 2018). The invaded sporozoites reside in clusters in the salivary glands till get a chance to invade the vertebrate host (Rosenberg, Wirtz et al. 1990; Ghosh and Jacobs-Lorena 2009). Though, studies targeting individual tissues such as midgut or salivary glands are valuable, but molecular mechanism of *Plasmodium* population alteration is yet to unravel (Simoes, Mlambo et al. 2017).

We hypothesize that for its survival *Plasmodium* must overcome at least two levels of competitive challenges **(Fig1)**. First one follows a 24-30 hrs pre-invasive phase of interaction initiated immediately after blood meal influencing: - (a) parasite development and adaptation to physiologically distinct but hostile gut environment than vertebrate host; (b) nutritional resources competition against exponentially proliferating gut microbes and (c) the barrier(s) infringement of gut epithelial prior maturation of peritrophic matrix, a unique but unresolved mechanism of self protection. A second phase follows post gut invasion of ookinetes which encompasses a direct interaction of (d) developing and maturing oocysts within midgut (8-10 days); (e) free circulatory sporozoites and hemocytes; and (f) salivary invaded sporozoites within salivary glands (10-16 days).

**Figure 1:**
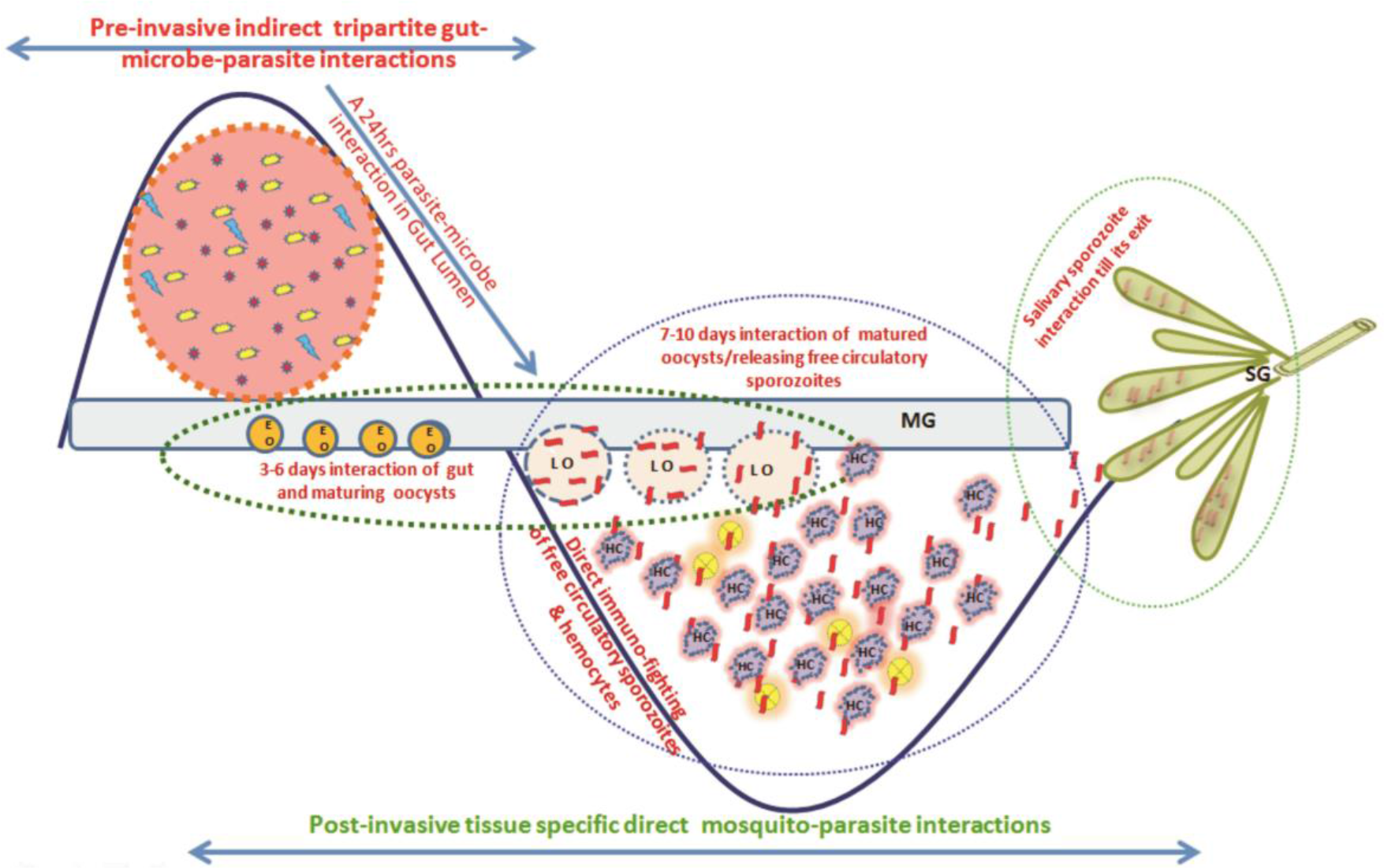
Proposed working hypothesis to decode a system-wide pre and post gut invasive phases of *P. vivax-*mosquito interactions: Immediately after infected blood meal, sexual developmental physiology of *Plasmodium* rapidly change to adapt mosquitoes’ hostile gut-lumen environment and progressively faces gut-flora boosted anti-*Plasmodium* immunity. Though the mechanism that how *Plasmodium* manages safe journey and survival from gut lumen ➙ gut epithelium ➙ hemolymph ➙ salivary gland ➙ vertebrate host is not fully known, but we propose and decode (i) a 24h-30hrs of **<u>pre-invasive phase of an indirect</u>** gut-microbe-parasite interaction in the gut lumen for ookinetes invasion; and (ii) a longer <u>**post gut invasive, a direct**</u> parasite-tissues such as Midgut (MG), hemocyte (HC) and salivary gland (SG) interactions, are crucial for the *Plasmodium* survival (Tevatiya et al., 2019; *BioRxiv*). Schematically, 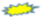 represents *Plasmodium* gametocytes; 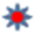, 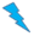 are different bacterial species, 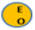 mustard yellow circle represents early gut invaded maturing oocysts (EO); 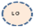 Peach circle with blue dotted boundary is Late rupturing oocysts (LO): 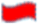 red ribbon is sporozoite, 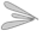 Salivary lobes, 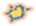 purple cloudy structure is hemocyte.

Thus, to decode the tissue specific molecular complexity/nature of interactions, we designed and carried out a system wide investigation. In the current report we demonstrate that how an early suppression of gut microbiome proliferation by *P. vivax* may support its survival during pre-invasive phase. While in second complementary report, we demonstrate that post gut invasion, a smart molecular relationship with individual tissues such as midgut, hemocytes, salivary glands, and strategic changes in the genetic makeup of *P. vivax* favors its survival in the mosquito host. (see Tevatiya et al., 2019; *BioRxiv*).

## Material and Methods

Technical overview presented in supplemental data Fig. S1

### Mosquito Rearing

*An. stephensi* colonies were reared in the central insectary facility at ICMR-National Institute of Malaria research (NIMR). A constant 28±2°C temperature and relative humidity of ~80% was maintained in the insectarium. Live rabbit was offered for blood meal for egg maturation and gonotrophic cycle maintenance (Sharma et al. 2015; Das De, Sharma et al. 2018).

### Metagenomic study

#### Tissue dissection and sample preparation

For the study *An. stephensi* pupae were reared in ethanol sterilized plastic cages fitted with autoclaved mesh cloth on the top. 10% sterile, fresh sugar solution was provided daily with a sterile cotton swab fitted in a test tube throughout the experiment. For metagenomics studies we collected the guts from 4-5 days old either sugar fed or blood fed adult female mosquitoes. Dissections were performed after surface sterilization of the mosquitoes using 75% ethanol for 1 minute in 20μl 1XSTE buffer and total gut DNA was extracted under aseptic conditions of the laminar air flow, as described earlier (Sharma, Sharma et al. 2014). In brief, the tissue was homogenized using hand held battery run homogenizer and volume was maintained to 50μl using STE and undesired protein was digested by treating with proteinase K. For DNA quality assessment ~5 μl of gDNA was loaded on 0.8% agarose gel to visualize the single intact band as the quality mark. The gel was run at standard conditions (Supplemental data Fig. S2). Quantification was performed using Qubit dsDNA BR kit (Thermo Fisher Scientific Inc.) after checking A_260/280_ ratio of 1 μl of each sample using Nanodrop 8000.

#### 16S rRNA based metagenomic sequencing and analysis

Using Nextera XT Index Kit (Illumina Inc.), the amplicon libraries were prepared from the qualified DNA samples. Primers were designed and synthesized using V3-V4 hyper-variable region of 16S rDNA gene (supplementary Table- ST1). The Illumina adaptors ligated amplicons were amplified by using i5 and i7 primers for multiplex indexing. Purification of the amplicon libraries was performed on 1X AMpureXP beads and checked for its quality with Bioanalyzer 2100 Agilent using a DNA1000 chip and quantification was done on fluorometer by Qubit dsDNA HS Assay kit (Life Technologies) (Supplemental data Fig. S3). A Paired-End (PE) sequencing was done with MiSeq technology and generated data was stitched into single end reads. Defined Operational Taxonomic Units (OTUs) were picked for taxonomic identity assignment using Quantitative Insights Into Microbial Ecology (QIIME version 1.9.1) and MEGAN softwares (Wang, Garrity et al. 2007; Price, Dehal et al. 2010).

#### Gut RNAseq analysis

Approximately one microgram purified total RNA from pooled 24-48hrs post blood fed 20 adult female mosquitoes guts, was subjected to double stranded cDNA library preparation (Clontech SMART^tm^) and sequencing (Illumina Technology), as described earlier (Zhu, Machleder et al. 2001). Briefly, purified ds cDNA sample (~200ng) was sheared using Covaris sonication method and the overhangs so generated were end-repaired before further processing. The paired end cDNA libraries were generated using illumina TruSeq Nano DNA HT Library Preparation Kit using 2×150 PE chemistry on NextSeq for generating ~1GB data as per described protocol. The end repaired fragments were subjected to enrichment by limited number of PCR cycles after adding a poly A-tail and adapter ligation. Library quantitation and qualification was performed using DNA high Sensitivity Assay Kit. The sequencing of whole transcriptomes, was performed on Illumina NextSeq. Trimmomatic v0.30 software was used to filter the raw reads. After removing adaptor sequences and low quality (QV < 20) reads, high quality clean reads were used to make *denovo* assembly using Trinity software (release r2013-02-25). CD-HIT-EST (Version 4.6) was used to remove the shorter redundant transcripts. All CDS were predicted from transcript using Transdecoder and selected longest frame transcripts were subjected for functional annotation using BLASTX against NR database and BLAST2GO program (see also Tevatiya et al., 2019; *BioRxiv*).

#### Artificial Membrane Feeding and *P. vivax* infection

The collection of the *P. vivax* infected patients’ blood samples was approved by the Ethics committee of NIMR, Delhi (ECR/NIMR/EC/2012/41). Prior collection of blood samples, a written informed consent (IC) was obtained from donors visiting to NIMR clinic. Venous blood was drawn into heparin-containing tubes and kept at 37°C till feeding. Overnight starved 4-5 days old female *An. stephensi* mosquitoes were fed using pre-optimized artificial membrane feeding assay (AMFA). Only full fed mosquitoes were maintained at optimal insectarium conditions and positive infection was confirmed by standard mercurochrome staining of gut oocysts readily observed under compound microscope. Desired tissue samples such as midgut, salivary glands, hemocytes were collected for subsequent analysis **(please see Fig.1 Panel A;**Tevatiya et al., 2019; *BioRxiv*). However, we excluded mosquito’s samples which showed poor/negative oocysts development in their gut.

#### RNA isolation and differential expression analysis

Total RNA was isolated from different tissues, originating from naïve, blood fed or *Plasmodium* infected *An*. *stephensi* mosquitoes (20 no.) and cDNA was synthesized using Verso cDNA synthesis kit (Thermo Fisher Scientific, #AB1453A) as per manufacturer protocol. Routine laboratory optimized RT PCR and agarose gel electrophoresis processes were followed for differential expression of the selected genes. Relative gene expression was performed by QuantiMix SYBR green dye (Thermo Scientific 2X DyNAmo Color Flash Sybr Green Master Mix Cal. No. F-416) in Eco-Real Time (Illumina, USA; Cat. No. EC-101-1001) or CFX-96 (Biorad, USA) Real-Time PCR machine. PCR cycle parameters included initial denaturation at 95°C for 15 min, followed by 44 cycles of 10 s at 95°C, 20 s at 55°C and 22 s at 72°C with a final extension of 15 s at 95°C, 15 s at 55°C and 15 s at 95°C. All qPCR measurements were performed in triplicates to rule out any possibility of biasness. At least three independent biological replicates were evaluated for better evaluation. Differential gene expression was evaluated using *ddCT* method and statistically analyzed by student *‘t’*test. List of primers presented in supplemental table (ST-2).

## Results

### Elizabethkingia and Pseudomonas predominate mosquito gut

To identify and catalogue gut associated bacteria, we sequenced and analysed a total of 3,68,138 Illumina raw reads originating from naive mosquito gut metagenomic library. Diversity richness indices such as Shannon-Weaver (1.662) and Simpson reciprocal (1.667) showed an optimal estimation and even distribution of species. A QIIME analysis at phylum level showed that mosquitoes gut dominantly harbours Bacteroidetes (73.13%); Proteobacteria (16.4%); Planctomycetes (4.10%); Firmicutes (2.3%), Verrucomicrobia (1.4%), Spirochaetes (1.2%), OD1 (0.4%) and 1.10% 16S reads remained unassigned **(Fig. 2a)**. At class level, the Flavobacteria *and* Gammaproteobacteria where *Elizabethkingia meningoseptica* and *Pseudomonas sp.* were most abundant gram negative bacteria, respectively. **(Fig. 2b)**

**Figure 2:**
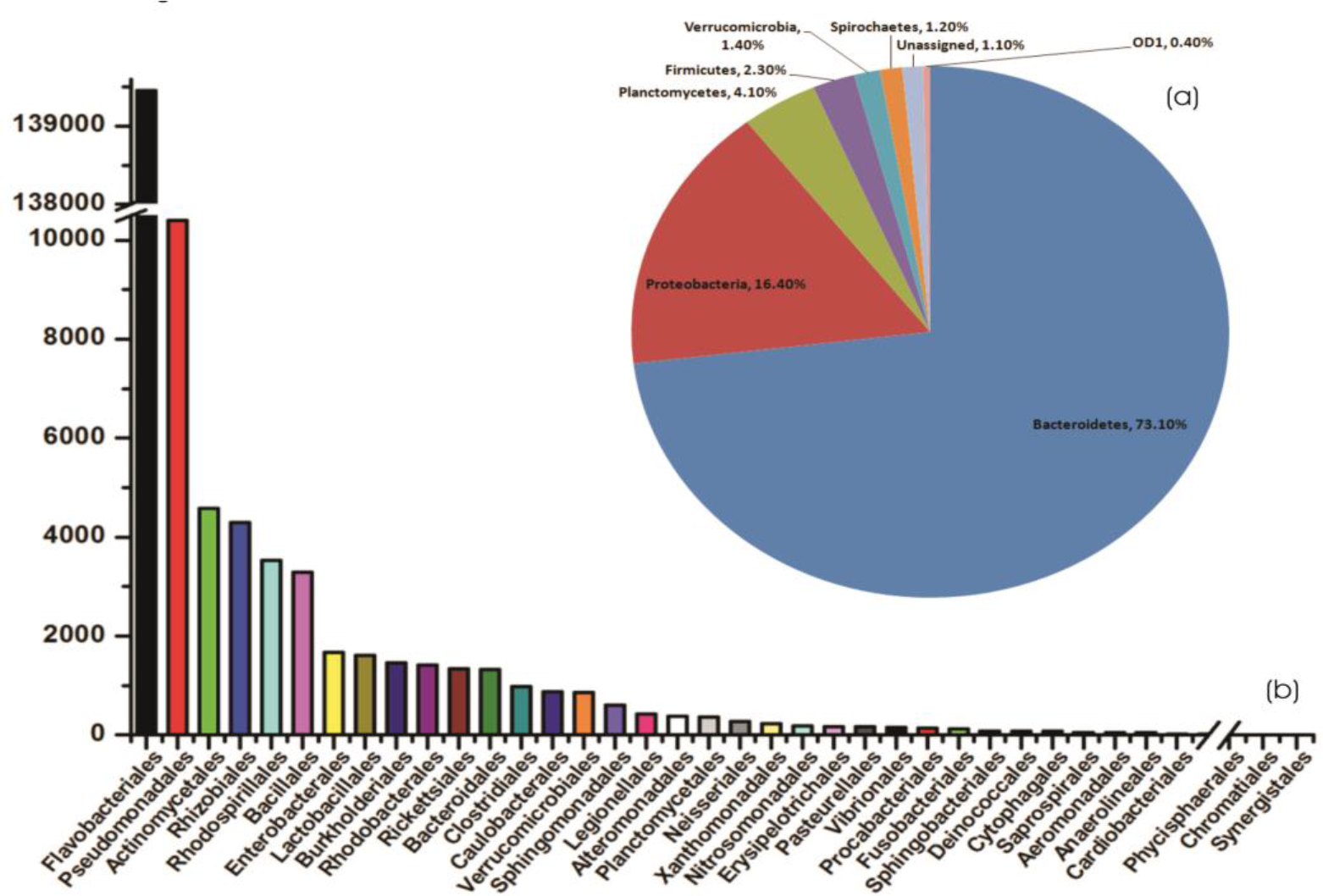
QIIME analysis based Gut microbial community structure in naïve mosquitoes: **(a)** microbial flora diversity of the mosquito gut, at phylum level showing dominant association of Bacteroidetes and Proteobacteria followed by Planctomycetes; Firmicutes, Verrucomicrobia, Spirochaetes; **(b)** The bar graph representing abundance of different bacteria at order level based on number of gut metagenomics reads, where *Flavobacteriales* and *Pseudomonadales* are the most abundant orders to which *Elizabethkingia* and *Pseusomonas* belongs respectively. Enterobacterales is also among top ten abundant orders which include Serratia.

### Blood meal alters the gut microbiome community structure

Supporting to previous studies, we also observed that blood meal gradually enriches gut total bacterial population till 24 hrs, which restore to their basal level within 48hrs of blood meal (Supplemental data Fig. S4). To further clarify that how blood meal influences individual bacterial population we catalogued and compared gut microbiome of naïve and 24hrs blood fed adult female mosquito guts. Alpha-diversity rarefaction curves estimate full extent of phylotype richness and quantifiable diversity estimation (Supplemental data Fig. S5). A normalized read count data comparison showed that blood meal not only enriches gut associated dominant Flavobacteria, but also favors modest enrichment of unique bacteria such as Bacillales, Lacto-bacilales, Spinghobacteriales, Rohocyclales **(Fig. 3a,b).**

**Figure 3:**
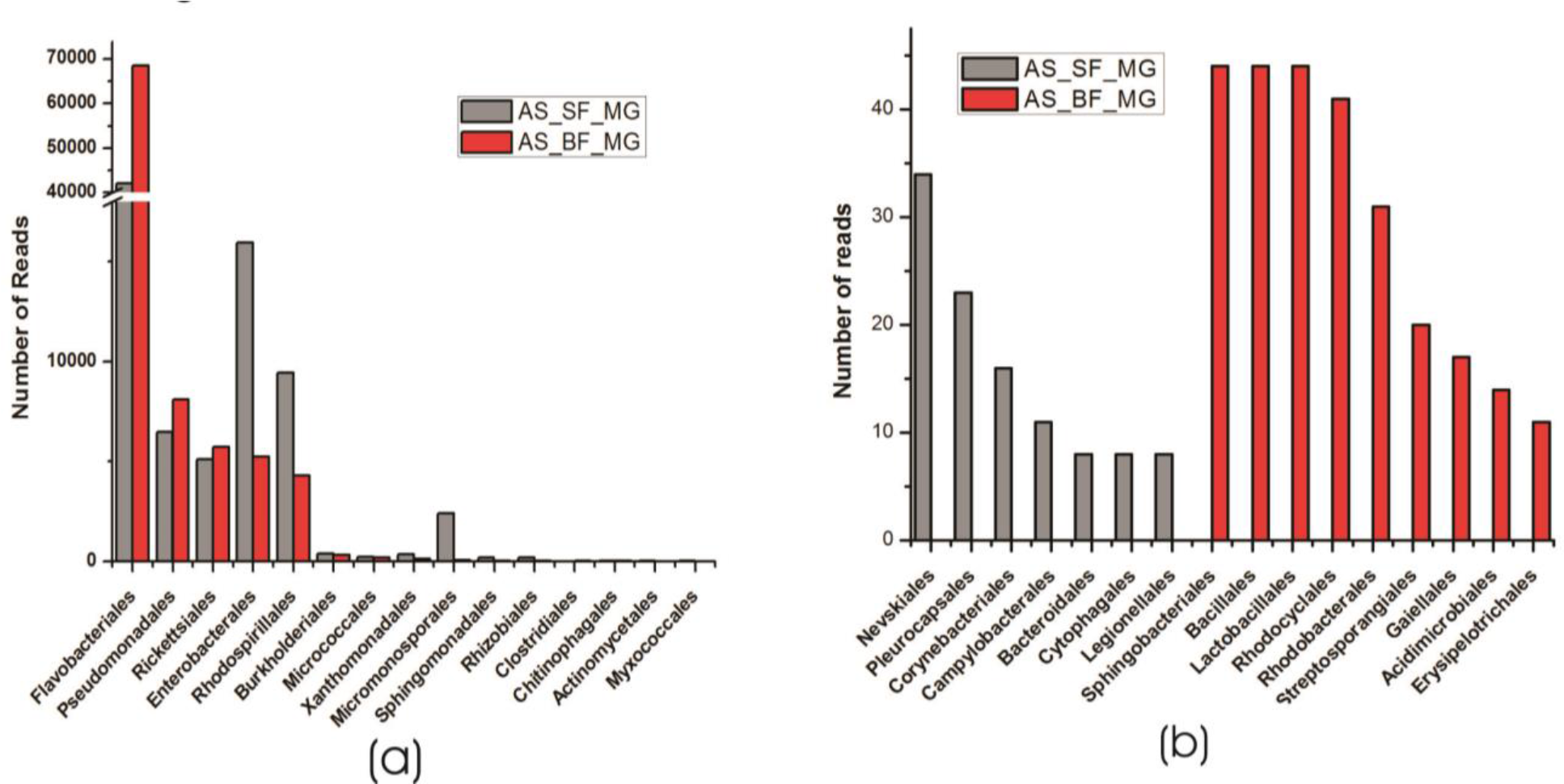
Blood meal alters gut community structure: **(a)** Comparative analysis of common bacterial community abundances among naïve and blood fed mosquitoes gut; **(b)** bar graph represents unique bacterial orders showing association with either sugar fed or blood fed mosquitoes gut.

To validate above observation, we examined relative abundances of selected bacterial species, by Real time PCR assay. We observed a relatively higher abundance of bacteria such as *Pseudomonas*, *Elizabethkingia* and *Serratia*, in the ovary and midgut than other tissues. However, within midgut, *Elizabethkingia* showed higher abundance than the *Pseudomonas* and *Serratia*, corroborating the metagenomic data (Fig. 4a, b,c,d; supplement table-ST3). Individual bacterial species such as *Elizabethkingia* (Flavobacteria), *Pseudomonas* and *Serratia* (Enterobacterace) also showed a gradual enrichment till 24 hours post blood feeding. However, post 30-hour blood meal digestion the bacterial population restored to the basal level of naïve mosquito midgut (Fig.5a,b,c).

**Figure 4:**
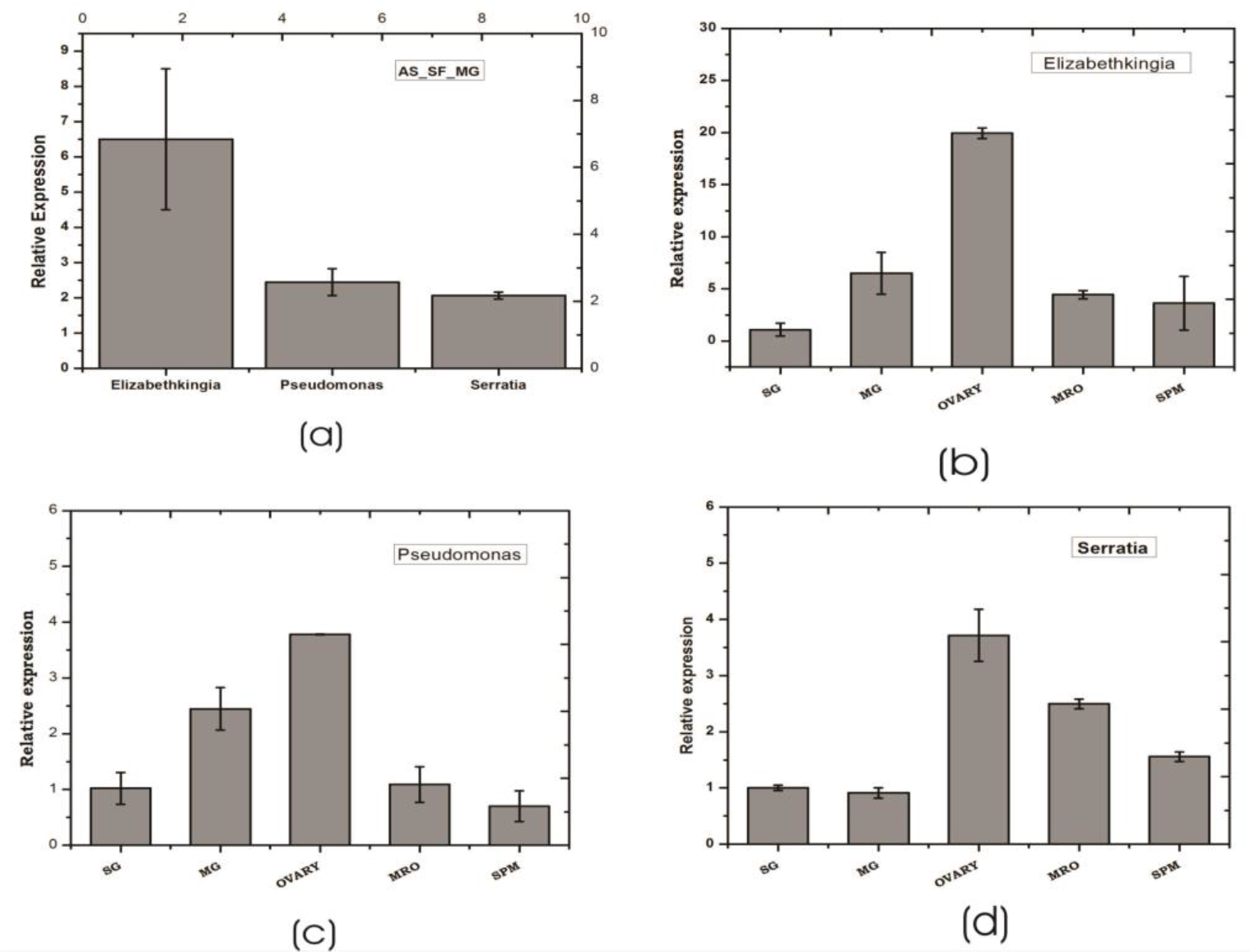
Tissue specific relative distribution of dominant endo-symbiotic bacteria in the naïve mosquitoes: Relative abundance of *Elizabethkingia, Pseudomonas, Serratia* in the naïve mosquito gut (a); tissue specific relative abundance of *Elizabethkingia* (b); *Pseudomonas* (c); and Serratia (d); SG: Salivary Gland, MG: midgut, MRO: Male reproductive organ, SPM: Spermathecae.

**Figure 5:**
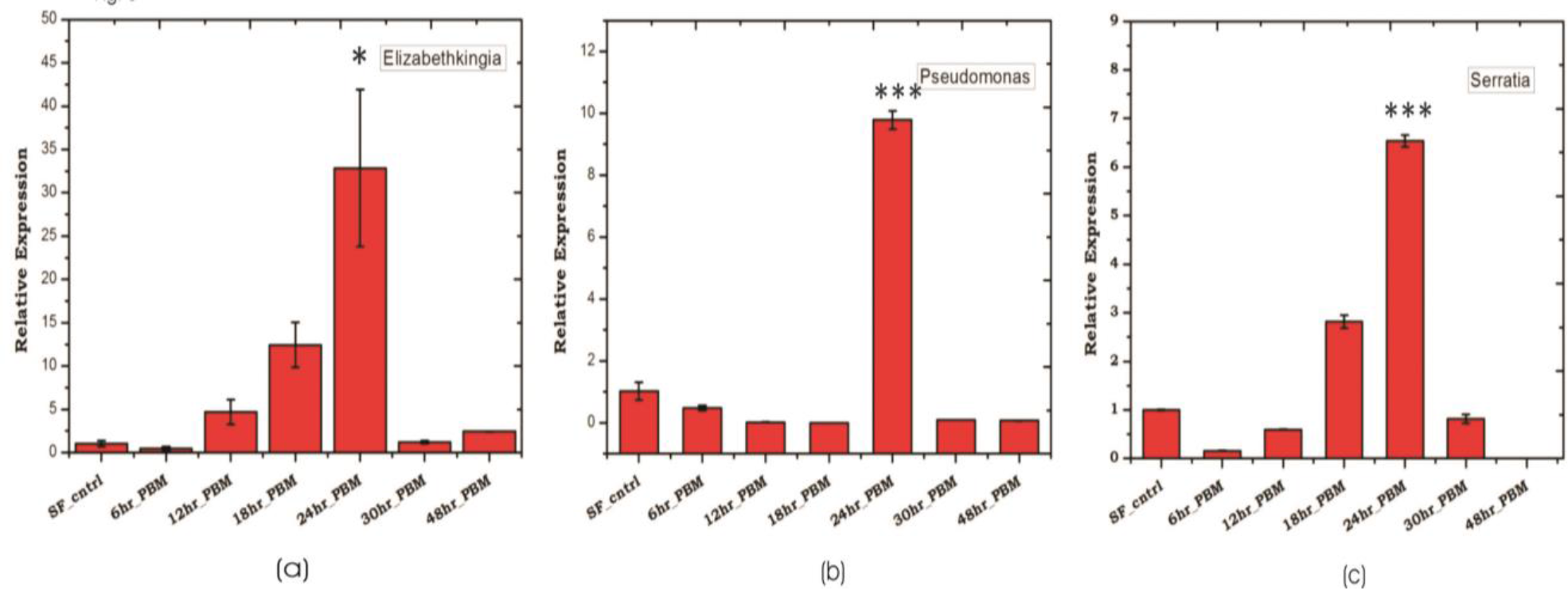
Blood feeding and species specific distribution of gut microbes: Time dependent relative abundance of **(a)** *Elizabethkingia* (p≤ 0.01); **(b)** *Pseudomonas* (p≤ 0.0005) and **(c)** *Serratia* (p≤ 0.001); in the blood fed mosquitoes gut. The gut tissue was collected at different time intervals of 6hr, 12hr, 18hr, 24hr, 30 hr and 48hr post blood feeding.

### RNAseq recovers molecular signatures of gut-microbe interaction

To establish a molecular/ functional relation of gut-microbe interaction, we analysed a total of 46,73,408 Illumina reads originating from 24h post blood fed gut RNAseq library (ST-4). Surprisingly, a species distribution analysis of 5,042 full length transcripts predicted that at least 90% transcripts sequences matched to insects, but ~10% transcripts i.e. 479 CDS showed a significant homology to microbial proteins (ST-5). Transcripts homolog to insects dominantly matched to *An. gambiae* (~72%), *An*. *sinensis* (~13%), *An*. *darlingi* (7~%), *Aedes aegypti* (~1.6%) and *Culex* (~1.1%) (Fig.6a). A close examination of BLASTx analysis of microbial sequences/transcripts further identified that at least 8% transcripts encodes proteins homologous to *Elizabethkingia* (EK) (Fig.6a), strengthening our finding that EK constitute a major gut endosymbiotic bacteria in *An. stephensi.* While remaining 2% transcripts showed a significant homology to other microbes such as *Annacalia alegera*; *Wolbachia* and viruses (ST-5).

**Figure 6:**
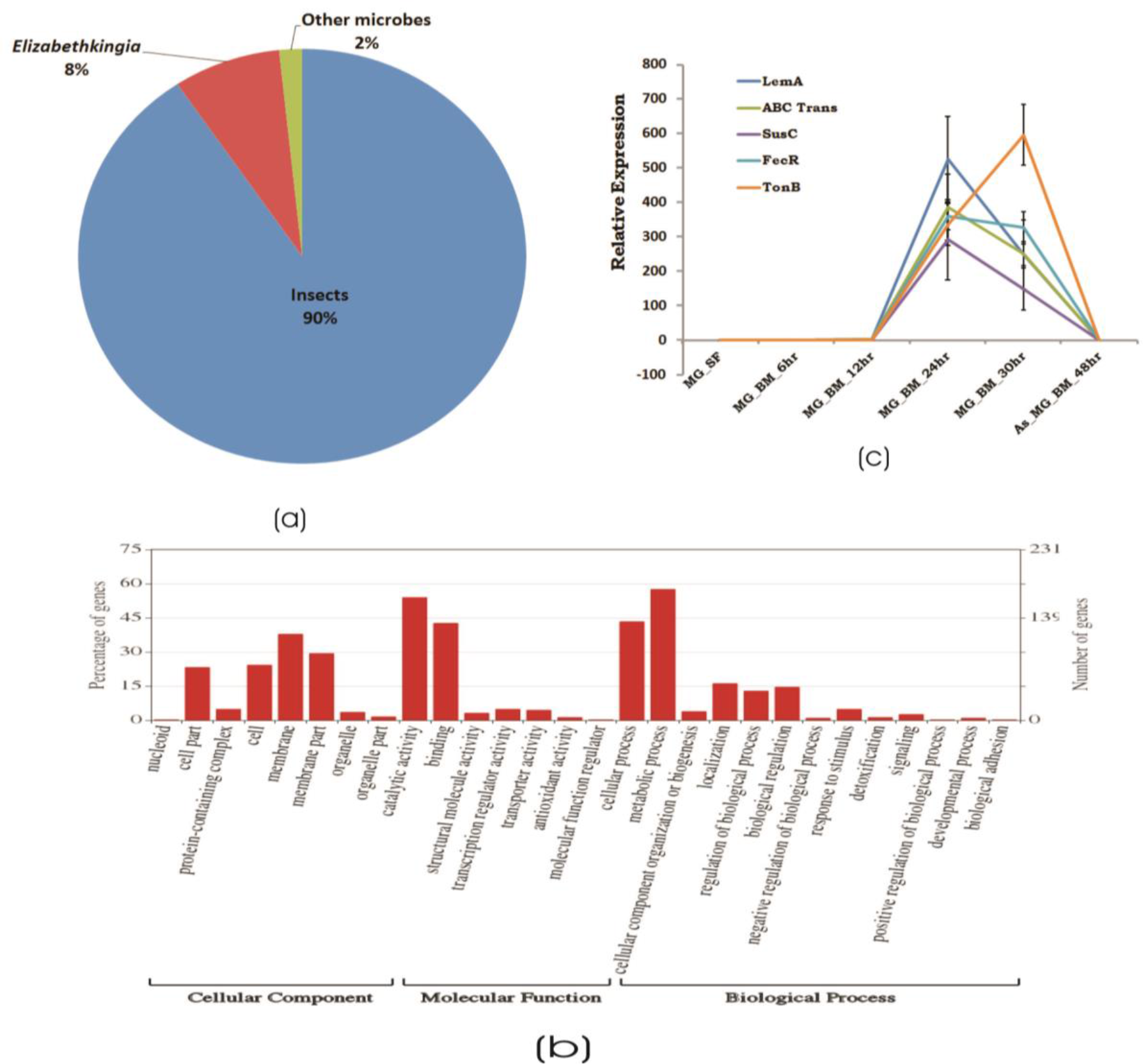
RNAseq identified microbial transcripts signatures: **(a)** Pie chart showing species distribution analysis of gut RNAseq data identifying transcripts having BLASTX homology to Insect (90%), *Elizabethkingia* as dominant gut endosymbiont bacteria (8%) and other microbes (2%); **(b)** molecular catalogue of identified EK transcripts; and **(c)** transcriptional profiling of bacterial EK specific transcripts in response to blood meal.

A comprehensive GO annotation of 391 putative transcripts indicated that EK bacterial species encodes diverse nature of proteins (Fig. 6b, ST-5). A transcriptional profiling of selected bacterial transcripts encoding LEM A, Ton-B dependent receptor, FecR, ABC transporter, SusC/Rag family protein showed enriched expression in response to blood feeding and digestion (Fig 6c/ST-6).

### 3. Early*Plasmodium vivax* Infection suppresses the gut microflora and immunity

We observed a significant loss in the gut bacterial population in *Plasmodium* infected mosquitoes which remained below detection limit till 36 hours (Fig. 7a). However, surprisingly, post 36hrs, the total bacterial population followed a gradual enrichment to multifold level till 10 days of gut infection (Fig 7a). Interestingly, EK and *Serratia* also showed a similar pattern of enrichment, except *Pseudomonas* whose population level remains least affected (Fig.7 b,c,d).

**Figure 7:**
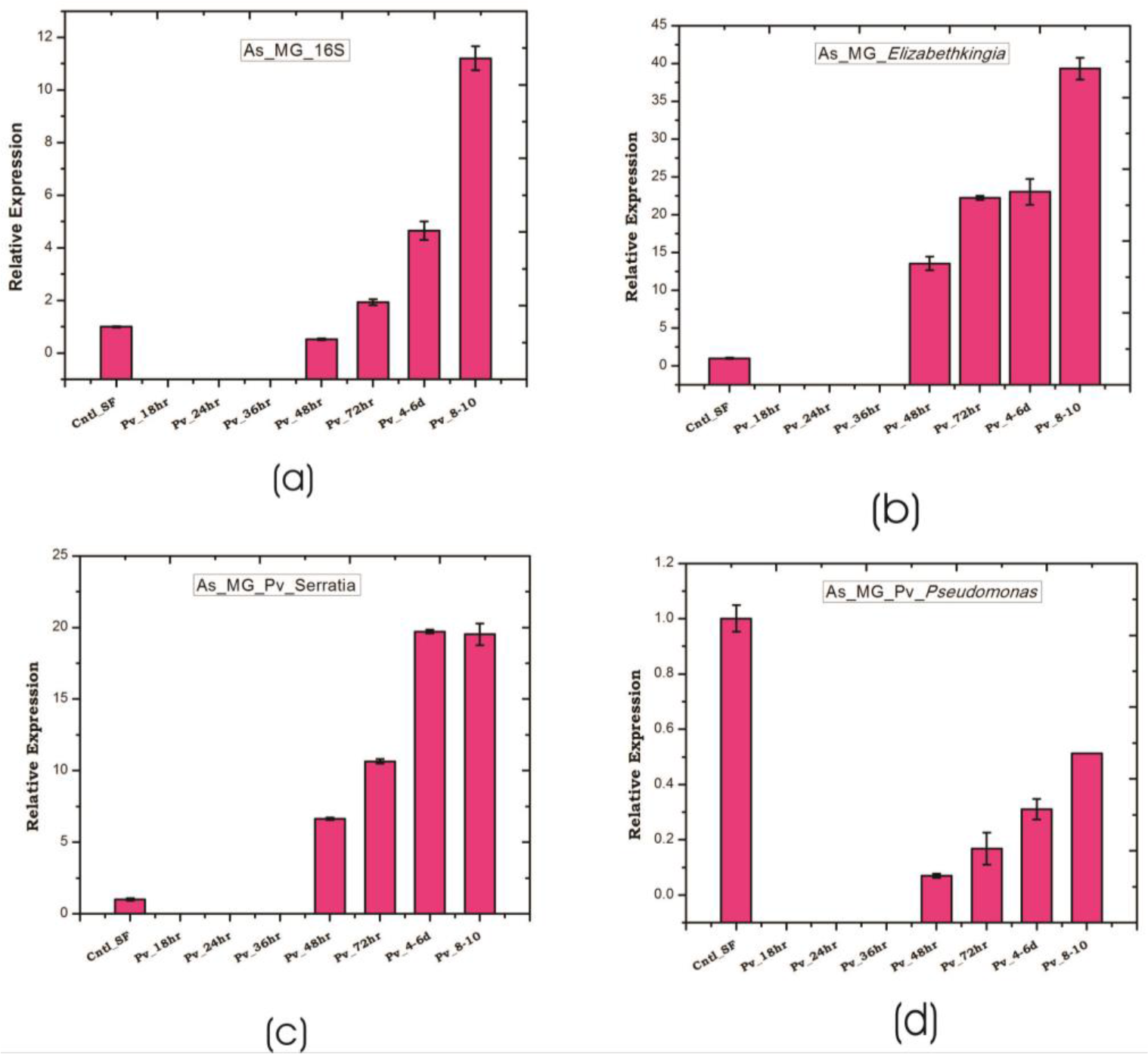
*P. vivax* infection cause early suppression and late restoration/enrichment of gut bacterial population: A time dependent relative quan.tification of gut microbiota in response to *Plasmodium vivax* infection showing enrichment 48h post infection (PI) of **(a)** *Total bacteria* (16S): p≤ 0.002/4-6DPI, p≤ 0.0004/8-10DPI**; (b)** *Elizabethkingia* p≤ 0.001/48hPI, p≤ 4.69E-05/72hPI, p≤ 0.001/4-6DPI, p≤ 0.0003/8-10DPI; **(c)** (Serratia p≤0.0001/48hPI, p≤ 8.73E-05//72hPI, p≤ 1.85E-05/4-6DPI, p≤ 0.0004)/ 8-10DPI; **(d)** *Pseudomonas*. DPI: days post infection.

Since, blood meal induced gut flora also boost gut immunity, we tested whether *P. vivax* infection influence gut immune response. A time dependent transcriptional profiling of all the selected anti-microbial peptides (also see Tevatiya et al., 2019; *BioRxiv*) showed a unique pattern of immune-suppression during pre-invasive phase of ookinetes to early oocysts development (Fig.8). All the tested immune transcripts showed expression enrichment only after 36 hours post infection. But, exceptionally, gambicin showed higher response than cecropin (C1, C2) and defensin (D1) (Fig. 8), suggesting its unique role against late oocysts development of *P. vivax*.

**Fig. 8.**
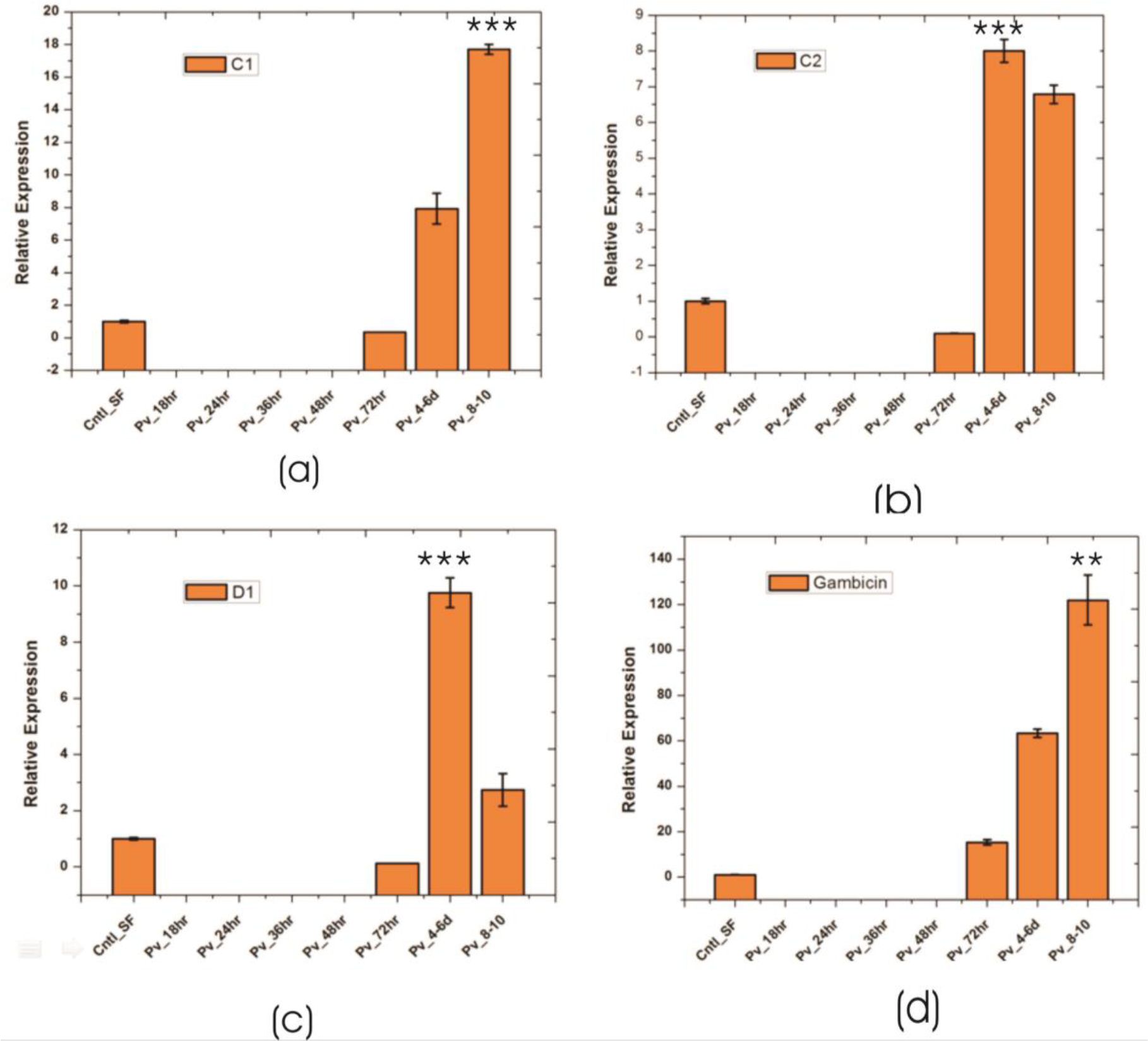
Relative quan.tification of gut immune transcripts in response to *Plasmodium vivax* infection: Transcriptional profiling of Antimicrobial peptides (AMPs) *C1* (*p≤ 9.00674E-05), C2 (p≤ 0.0005), D1 (p≤ 0.0008), Gambicin (p≤ 0.002)* showing early suppression of gut immunity which restored after three days of *P. vivax* infected blood meal. C1-C2: cecropin1 and cecropin2; D1: defensin1.

### Laboratory reared *Anopheles stephensi* harbour *Wolbachia bacteria*

Surprisingly, a qualified subset of 250bp long metgenomic sequencing reads (6532 blood fed and 6154 naïve mosquitoes gut) showed 100% identity to *Wolbachia* endosymbiont of *Chrysomya megacephala* (Accession #CP021120.1; Supplemental data S6a/also see Supplemental *FASTA file* S8). Also identification of at least 7 mRNA transcripts, originating from distinct gut RNAseq libraries and encoding different *Wolbachia* homolog proteins (ST-7/S6b), further predicts novel Wolbachia association. An ongoing similar comparative gut metagenomic analysis of Indian vector *An. culicifacies*, (unpublished) reared in same insectarium environment, did not yielded a single sequence of *Wolbachia* origin, supporting that *An. stephensi* may exclusively harbour novel *Wolbachia* bacterial species.

## Discussion

Using meta-transcriptomic strategy, we targeted to decode tripartite gut-microbe-*P. vivax* interaction in the mosquito host *An. stephensi*. Our metagenomic study identifies *Elizabethkingia* and *Pseudomonas*, as dominant gut inhabiting bacteria in the laboratory reared naïve adult female mosquitoes. In response to blood meal, we observed a significant alteration of gut microbial community structure and enrichment of dominant bacterial species e.g. *Elizabethkingia sp*. (Falvibacteriace), *Pseudomonas* (Pseudomonadales), and *Serratia* (Enterobacteriace). Previous several studies have also reported a similar pattern of gut microbe enrichment (Tchioffo, Boissiere et al. 2015; Muturi, Dunlap et al. 2019), but nature of gut-microbe interactions, especially microbial proteins facilitating blood meal digestion, remains poorly known (Chen, Blom et al. 2017). Available draft genome sequence of cultured bacterial species predicts several metabolic pathways, but no functional relation has been established (Kukutla, Lindberg et al. 2013; Pei, Hill-Clemons et al. 2015).

A functional annotation of at least ~391 *Elizabethkingia* transcripts identified from blood fed mosquitoes gut-RNAseq data provide a direct evidence of ‘*in vivo*’ metabolically active proteins having role in blood meal digestion. Till 30hrs of post blood meal, an enriched expression of transcripts such as LEM-A, Ton-B dependent receptor, FecR, ABC transporter suggested their important role in iron-metabolism. Possibly this is accomplished through siderophore uptake and oxidative stress management, a mechanism benefiting mosquito’s survival and reproductive outcome (Koster 2001; Mettrick and Lamont 2009; Wang, Xu et al. 2016).

It is known that gut endosymbionts also serve as potent modulators of sexual development and transmission of the malaria parasite in *Anopheles* mosquitoes (Weiss and Aksoy 2011; Tchioffo, Boissiere et al. 2013). This antagonistic relationship of gut bacteria has been observed in the sporogonic development of *Plasmodium* in several Anopheline mosquitoes (Pumpuni, Beier et al. 1993; Dong, Manfredini et al. 2009; Tchioffo, Boissiere et al. 2013). Introduction of *E. coli*, *Pseudomonas* and *Serratia* by oral feeding reduces the gut oocyst load in *An. gambiae* (Tchioffo, Boissiere et al. 2013), but species specific interaction of the *Plasmodium* and bacteria remains un-clarified. In our infectivity assay we observed that *P. vivax* disables bacterial proliferation to keep an immuno-suppression till invasion to gut epithelium.

Though, it is unknown that how sexual stages of *Plasmodium* utilize ingested blood iron in the mosquito gut, but earlier study suggests that an iron depleted blood inhibits *P. falciparum* gametocyte activation, and hence the infectivity (Ke, Sigala et al. 2014). Thus, we hypothesize first 24 hrs of gut-microbe-*Plasmodium* interaction in the gut lumen are crucial for *Plasmodium* survival, where it may limit the availability of iron/nutrient required for bacterial growth (Clark, Goheen et al. 2014). Corroborating to earlier studies, we also observed that mosquitoes were able to restore basal level of gut flora within 30hrs of uninfected blood meal digestion (Das De et al., 2018). However, surprisingly, *P. vivax* infection caused a major shift in gut flora restoration to an enriched state after 48 hrs in *P. vivax* infection. Interestingly, this shift of bacterial enrichment boosted a similar a pattern of gut immunity induction, till late oocysts exited gut epithelium (Fig.8). Together, we hypothesize that in the gut lumen, gut-microbe-*P. vivax* interaction undergoes a unique ‘flip-show’ where an early suppression of gut bacteria may favor *Plasmodium* survival, but late phase gut immunity activation may restrict gut oocysts population. A late phase anti-*Plasmodium* immunity has also been suggested in other mosquito-Parasite interaction studies (Clayton, Dong et al. 2014). Since, we observed this pattern repeatedly for at least four independent experiments, thus it is very unlikely that it may be an undisclosed confounding effect of a blood sample originating from patient having antibiotic treatment before diagnosis (Gendrin, Rodgers et al. 2015).

Paratransgenesis approaches for manipulating gut endosymbionts such as *Elizabethkingia*, *Serratia* to block parasite development are under progress (Chen, Bagdasarian et al. 2015; Wang, Dos-Santos et al. 2017; Koosha, Vatandoost et al. 2018). A dominant association of tested *Elizabethkingia*, *Pseudomonas*, *Serratia* with mosquito ovaries/eggs, and subsequent validation of trans ovarian transfer from F1, F2 and F3 generation (See Supplementary data S7), supports an idea to select and target them for future manipulation Alternatively, manipulating intracellular endosymbiont such as *Wolbachia* induced male sterility and pathogen development inhibition, are rapidly gaining much attention for vector borne disease control program (Bourtzis, Dobson et al. 2014; Jayakrishnan, Sudhikumar et al. 2018). Trial releases of *Wolbachia* inhabiting mosquitoes now being proved as a tool to reduce dengue cases in several countries (Dorigatti, McCormack et al. 2018; Garcia, Sylvestre et al. 2019). A laboratory validation of similar strategy in *Anopheline* mosquitoes for malaria control is also in progress (Bian, Joshi et al. 2013).

A surprising finding of at least ~6% metagenomic sequences, and *Wolbachia* homolog protein encoding transcripts, further established a natural association of a novel *Wolbachia* bacteria in laboratory reared mosquitoes. Thus, we believe a systemic evaluation and validation of *Wolbachia* interaction influencing *Plasmodium* development, and cytoplasmic incompatibility in *An. stephensi*, may be valuable to design novel tool to fight malaria in India.

## Conclusion

Several studies prove that immediately after blood feeding, a vital tripartite interaction occurs among mosquito-microbe-parasite in the mosquito’s gut lumen. But the molecular basis that how *Plasmodium* manages its survival, development and transmission is not well known. For the first time we establish that *P. vivax* causes an early suppression of gut microbial population, possibly by altering iron metabolism and nutritional physiology. And by this strategy parasite not only weakens gut immunity, but also favors successful invasion and development in the mosquito *Anopheles stephensi* (Fig. 9). With current data we further propose that late oocysts/brusting sporozoites alters gut bacterial susceptibility to boost late phase immunity, a plausible mechanism to restrict the *Plasmodium* population (see *Tevatiya et al., 2019*; *BioRxiv*)

**Figure 9:**
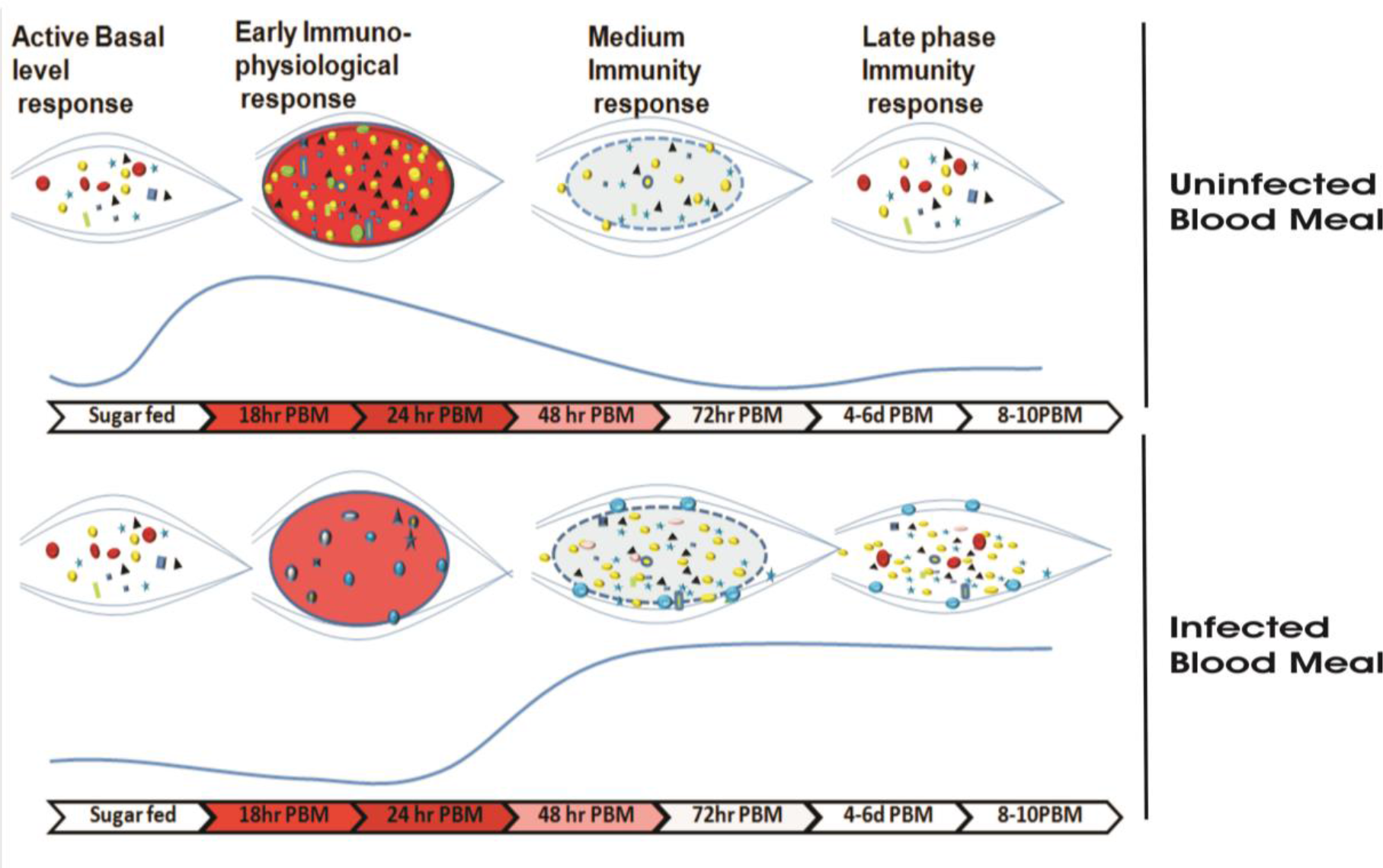
Survival strategy model of *P. vivax* during pre-invasive phase of development inside the mosquito gut: A schematic representation of the microbial distribution in response to blood feeding versus *Plasmodium* infected blood meal. In absence of *Plasmodium*, after normal blood feeding rapid proliferation of gut bacteria occurs in a nutrient rich medium (of which Fe is indispensable to bacterial growth) during a time window of 18-24 hr. Mosquito induces innate immune response (via production of various AMPs) to this incremental bacterial growth in order to tame the bacterial load and restore it to a basal level by 48 hrs. In presence of *Plasmodium*, *Plasmodium vivax* suppresses the proliferation of gut bacteria possibly by altering Fe metabolism or nutrition physiology during initial hours (18hr-24hr) in a bid to dampen the mosquito innate immune response and to shore up ookinete invasion. In a direct competition for nutritional resources within the gut lumen between the parasite and bacteria, parasite overcomes the bacteria. Parasite leaves the lumen and encysts beneath the basal lamina. After 48hr post blood meal, in absence of competition with the parasite in gut lumen allows the bacteria to proliferate possibly by feeding on undigested food left in lumen. As the bacterial load rises, it leads to activation of mosquito innate immune responses followed by synthesis of various AMPs which not only limit the bacterial load but also limit the medium and late oocyst development. 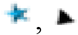, 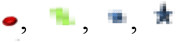, represent different bacteria residing the gut 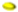, *Elizabethkingia*, 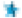 *Pseudomonas*, 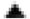 *Serratia*; 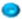 *Plasmodium viva;*. 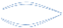 *Midgut;* 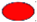 Blood bolus after normal blood feeding, 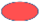 blood bolus after *Plasmodium* infected blood uptake. 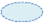. Peritrophic matrix after blood digestion.

## Supporting information

Comparative_Metagenomic

NR_Annotation

Fasta File

## Data Submission Detail

The sequence data has been submitted to NCBI SRA database under following accession number: SAMN10496496- DNA_AS_SF_MG; SAMN10439711-AS_MG_BF_DNA for individual samples of metagenomics data described in the manuscript. The sequences of Blood fed midgut RNASeq data is submitted with accession number-SRR8580010. All other data is included as Supplementary material.

## Acknowledgement

Work in the laboratory is supported by Indian Council of Medical Research (ICMR), Government of India (3/1/3PDF(13)/2016-HRD). RKD is a recipient of a DBT sponsored Ramalingaswami Fellowship. Authors thanks to NIMR-clinical facility support. Authors also thank ICMR-NIMR for basic infrastructure and instrumentation support. We thank Kunwarjeet Singh for technical assistance and mosquito rearing. We are thankful to Xcelris Labs Limited Ahmedabad, Gujarat, India, and NGB Patparganj, Delhi for the meta-transcriptomic sequencing services.

## Authors’ contribution

PS, RKD,KCP idea and hypothesis generation, conceived and designed the experiments; PS, CC, SK, JR, ST, TDD contributed to design and performing the experiments, data acquisition, writing and editing; PS, KCP, RKD data analysis and interpretation, data presentation, contributed reagents/ materials/Analysis tools, wrote, reviewed, edited, and finalized MS. All authors read and approved the final manuscript.

## References

Attardo, G. M., I. A. Hansen, et al. (2005). “Nutritional regulation of vitellogenesis in mosquitoes: implications for anautogeny.” Insect Biochem Mol Biol 35(7): 661–675.

Belachew, E. B. (2018). “Immune Response and Evasion Mechanisms of Plasmodium falciparum Parasites.” J Immunol Res 2018: 6529681.

Bennink, S., M. J. Kiesow, et al. (2016). “The development of malaria parasites in the mosquito midgut.” Cell Microbiol 18(7): 905–918.

Bian, G., D. Joshi, et al. (2013). “Wolbachia invades Anopheles stephensi populations and induces refractoriness to Plasmodium infection.” Science 340(6133): 748–751.

Bourtzis, K., S. L. Dobson, et al. (2014). “Harnessing mosquito-Wolbachia symbiosis for vector and disease control.” Acta Trop 132 Suppl: S150–163.

Chen, S., M. Bagdasarian, et al. (2015). “Elizabethkingia anophelis: molecular manipulation and interactions with mosquito hosts.” Appl Environ Microbiol 81(6): 2233–2243.

Chen, S., J. Blom, et al. (2017). “Genomic, Physiologic, and Symbiotic Characterization of Serratia marcescens Strains Isolated from the Mosquito Anopheles stephensi.” Front Microbiol 8: 1483.

Clark, M. A., M. M. Goheen, et al. (2014). “Influence of host iron status on Plasmodium falciparum infection.” Front Pharmacol 5: 84.

Clayton, A. M., Y. Dong, et al. (2014). “The Anopheles innate immune system in the defense against malaria infection.” J Innate Immun 6(2): 169–181.

Das De, T., P. Sharma, et al. (2018). “Interorgan Molecular Communication Strategies of “Local” and “Systemic” Innate Immune Responses in Mosquito Anopheles stephensi.” Front Immunol 9: 148.

Dixit, R., M. Rawat, et al. (2011). “Salivary gland transcriptome analysis in response to sugar feeding in malaria vector Anopheles stephensi.” J Insect Physiol 57(10): 1399–1406.

Dong, Y., F. Manfredini, et al. (2009). “Implication of the mosquito midgut microbiota in the defense against malaria parasites.” PLoS Pathog 5(5): e1000423.

Dorigatti, I., C. McCormack, et al. (2018). “Using Wolbachia for Dengue Control: Insights from Modelling.” Trends Parasitol 34(2): 102–113.

Drexler, A. L., Y. Vodovotz, et al. (2008). “Plasmodium development in the mosquito: biology bottlenecks and opportunities for mathematical modeling.” Trends Parasitol 24(8): 333–336.

Gaio Ade, O., D. S. Gusmao, et al. (2011). “Contribution of midgut bacteria to blood digestion and egg production in aedes aegypti (diptera: culicidae) (L.).” Parasit Vectors 4: 105.

Garcia, G. A., G. Sylvestre, et al. (2019). “Matching the genetics of released and local Aedes aegypti populations is critical to assure Wolbachia invasion.” PLoS Negl Trop Dis 13(1): e0007023.

Gendrin, M., F. H. Rodgers, et al. (2015). “Antibiotics in ingested human blood affect the mosquito microbiota and capacity to transmit malaria.” Nat Commun 6: 5921.

Ghosh, A. K. and M. Jacobs-Lorena (2009). “Plasmodium sporozoite invasion of the mosquito salivary gland.” Curr Opin Microbiol 12(4): 394–400.

Jayakrishnan, L., A. V. Sudhikumar, et al. (2018). “Role of gut inhabitants on vectorial capacity of mosquitoes.” J Vector Borne Dis 55(2): 69–78.

Ke, H., P. A. Sigala, et al. (2014). “The heme biosynthesis pathway is essential for Plasmodium falciparum development in mosquito stage but not in blood stages.” J Biol Chem 289(50): 34827–34837.

Koosha, M., H. Vatandoost, et al. (2018). “Delivery of a Genetically Marked Serratia AS1 to Medically Important Arthropods for Use in RNAi and Paratransgenic Control Strategies.” Microb Ecol.

Koster, W. (2001). “ABC transporter-mediated uptake of iron, siderophores, heme and vitamin B12.” Res Microbiol 152(3-4): 291–301.

Kukutla, P., B. G. Lindberg, et al. (2013). “Draft Genome Sequences of Elizabethkingia anophelis Strains R26T and Ag1 from the Midgut of the Malaria Mosquito Anopheles gambiae.” Genome Announc 1(6).

Mettrick, K. A. and I. L. Lamont (2009). “Different roles for anti-sigma factors in siderophore signalling pathways of Pseudomonas aeruginosa.” Mol Microbiol 74(5): 1257–1271.

Muturi, E. J., C. Dunlap, et al. (2019). “Host blood-meal source has a strong impact on gut microbiota of Aedes aegypti.” FEMS Microbiol Ecol 95(1).

Noden, B. H., J. A. Vaughan, et al. (2011). “Mosquito ingestion of antibodies against mosquito midgut microbiota improves conversion of ookinetes to oocysts for Plasmodium falciparum, but not P. yoelii.” Parasitol Int 60(4): 440–446.

O’Neill, S. L., R. Giordano, et al. (1992). “16S rRNA phylogenetic analysis of the bacterial endosymbionts associated with cytoplasmic incompatibility in insects.” Proc Natl Acad Sci U S A 89(7): 2699–2702.

Pei, D., C. Hill-Clemons, et al. (2015). “Draft Genome Sequences of Two Strains of Serratia spp. from the Midgut of the Malaria Mosquito Anopheles gambiae.” Genome Announc 3(2).

Price, M. N., P. S. Dehal, et al. (2010). “FastTree 2--approximately maximum-likelihood trees for large alignments.” PLoS One 5(3): e9490.

Pumpuni, C. B., M. S. Beier, et al. (1993). “Plasmodium falciparum: inhibition of sporogonic development in Anopheles stephensi by gram-negative bacteria.” Exp Parasitol 77(2): 195–199.

Pumpuni, C. B., J. Demaio, et al. (1996). “Bacterial population dynamics in three anopheline species: the impact on Plasmodium sporogonic development.” Am J Trop Med Hyg 54(2): 214–218.

Richards, S. L., S. L. Anderson, et al. (2012). “Effects of blood meal source on the reproduction of Culex pipiens quinquefasciatus (Diptera: Culicidae).” J Vector Ecol 37(1): 1–7.

Rodgers, F. H., M. Gendrin, et al. (2017). “Microbiota-induced peritrophic matrix regulates midgut homeostasis and prevents systemic infection of malaria vector mosquitoes.” PLoS Pathog 13(5): e1006391.

Romoli, O. and M. Gendrin (2018). “The tripartite interactions between the mosquito, its microbiota and Plasmodium.” Parasit Vectors 11(1): 200.

Rosenberg, R., R. A. Wirtz, et al. (1990). “An estimation of the number of malaria sporozoites ejected by a feeding mosquito.” Trans R Soc Trop Med Hyg 84(2): 209–212.

Sharma, P., S. Sharma, et al. (2015). “Unraveling dual feeding associated molecular complexity of salivary glands in the mosquito Anopheles culicifacies.” Biol Open 4(8): 1002–1015.

Simoes, M. L., G. Mlambo, et al. (2017). “Immune Regulation of Plasmodium Is Anopheles Species Specific and Infection Intensity Dependent.” MBio 8(5).

Simonetti, A. B. (1996). “The biology of malarial parasite in the mosquito--a review.” Mem Inst Oswaldo Cruz 91(5): 519–541.

Smith, R. C., J. Vega-Rodriguez, et al. (2014). “The Plasmodium bottleneck: malaria parasite losses in the mosquito vector.” Mem Inst Oswaldo Cruz 109(5): 644–661.

Tchioffo, M. T., A. Boissiere, et al. (2015). “Dynamics of Bacterial Community Composition in the Malaria Mosquito’s Epithelia.” Front Microbiol 6: 1500.

Tchioffo, M. T., A. Boissiere, et al. (2013). “Modulation of malaria infection in Anopheles gambiae mosquitoes exposed to natural midgut bacteria.” PLoS One 8(12): e81663.

Wang, Q., G. M. Garrity, et al. (2007). “Naive Bayesian classifier for rapid assignment of rRNA sequences into the new bacterial taxonomy.” Appl Environ Microbiol 73(16): 5261–5267.

Wang, R., H. Xu, et al. (2016). “A TonB-dependent receptor regulates antifungal HSAF biosynthesis in Lysobacter.” Scientific Reports 6: 26881.

Wang, S., A. L. A. Dos-Santos, et al. (2017). “Driving mosquito refractoriness to Plasmodium falciparum with engineered symbiotic bacteria.” Science 357(6358): 1399–1402.

Weiss, B. and S. Aksoy (2011). “Microbiome influences on insect host vector competence.” Trends Parasitol 27(11): 514–522.

Zhu, Y. Y., E. M. Machleder, et al. (2001). “Reverse transcriptase template switching: a SMART approach for full-length cDNA library construction.” Biotechniques 30(4): 892–897.

