## Supplemental-Data_File_S1-S7 for "Altered gut microbiota and immunity defines *Plasmodium vivax* survival in *Anopheles stephensi*"

Fig:S1

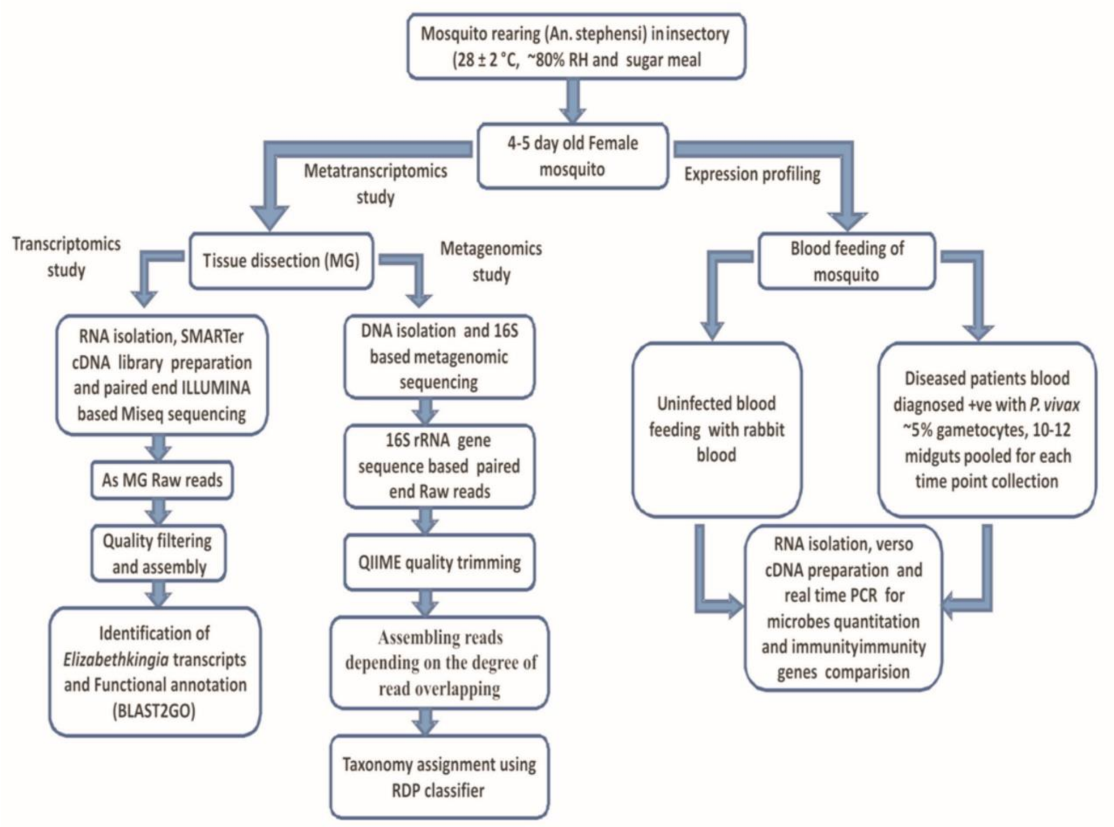


**Fig.S1 Technical work plan to decode molecular complexity of mosquito-microbiota-parasite interaction.** In this study the two way approach including the metatranscriptomics based bioinformatics study and the Real time based relative abundance of the selected bacteria and related genes is followed to find the role of the residing bacteria in blood feeding and parasite transmission.

Fig. S2

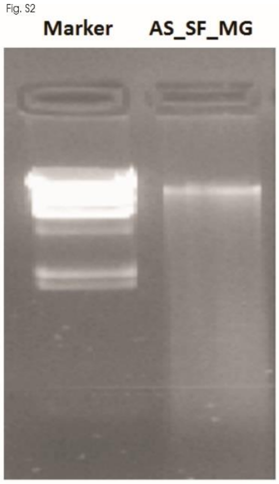


**S2:** Agarose Gel Picture showing the quality check of gDNA of the sugar fed *Anopheles stephensi* midgut. First lane is the marker and 2nd lane is of AS_SF_MG.

Fig. S3

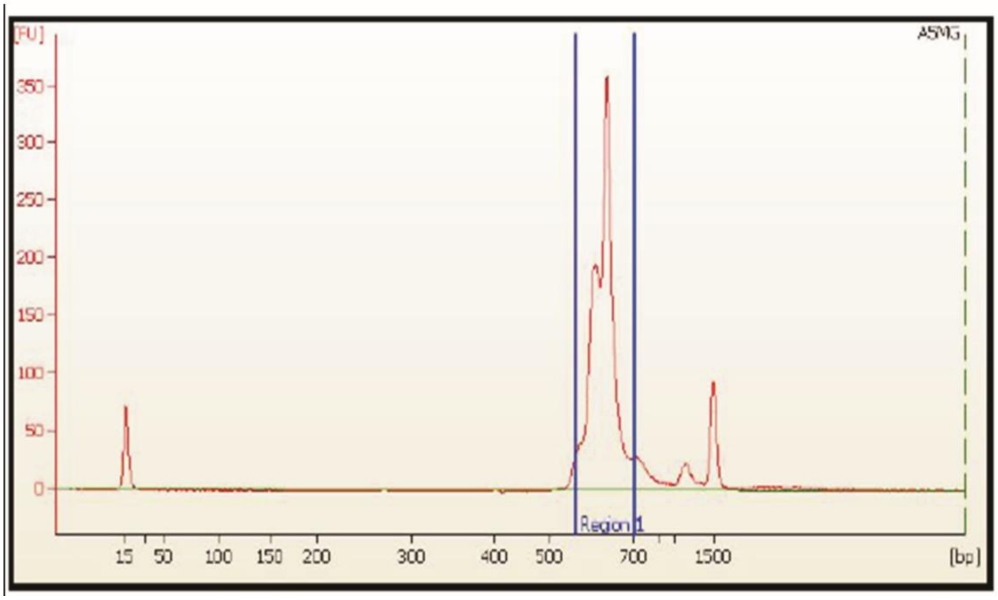


**Fig. S3**: The electropherogram of the Bioanalyzer 2100 analysis for the amplicon library which was purified by 1X AMpureXP beads showing the major peak within 500bp and 700 bp that was taken for subsequent bacterial sequencing, profiling and analysis.

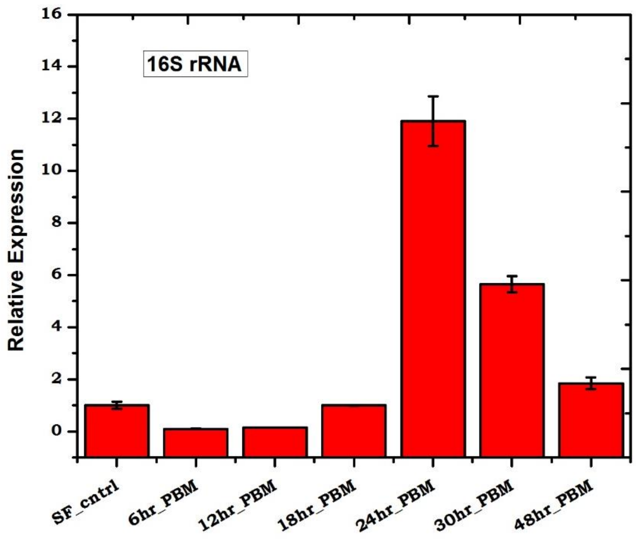


**Fig. S4:** Real-time PCR based estimation of relative abundance of gut bacterial population in response to blood feeding. The figure shows the highest 16S rRNA expression post 24 hrs of blood feeding as compared to sugar feeding and early blood fed stages. At 48hrs PBM abundance of the bacteria retains at a level comparable to its sugar fed

(a)

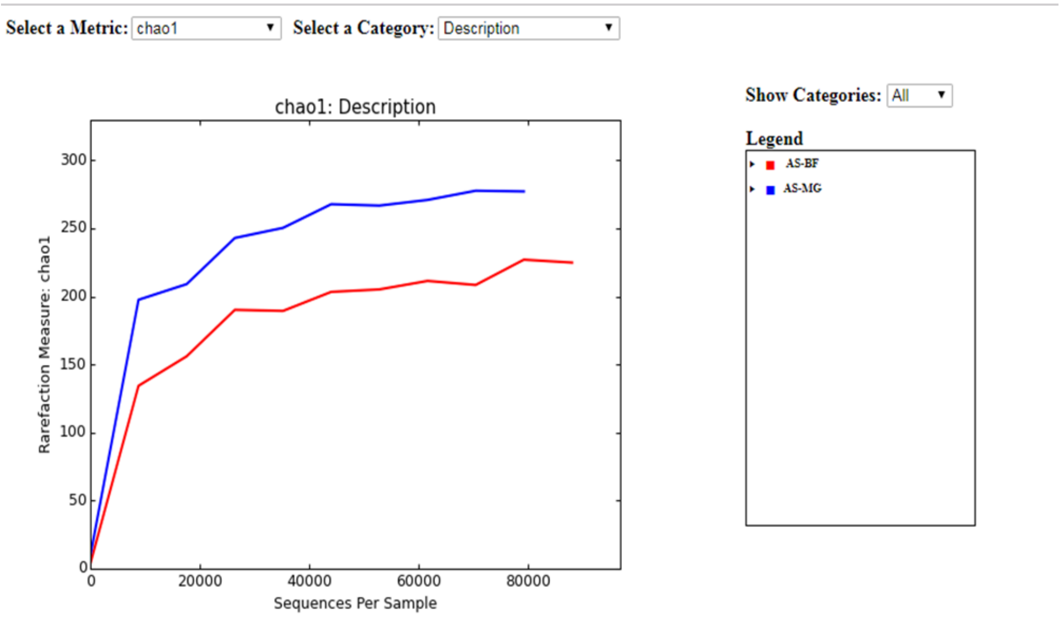


(b)

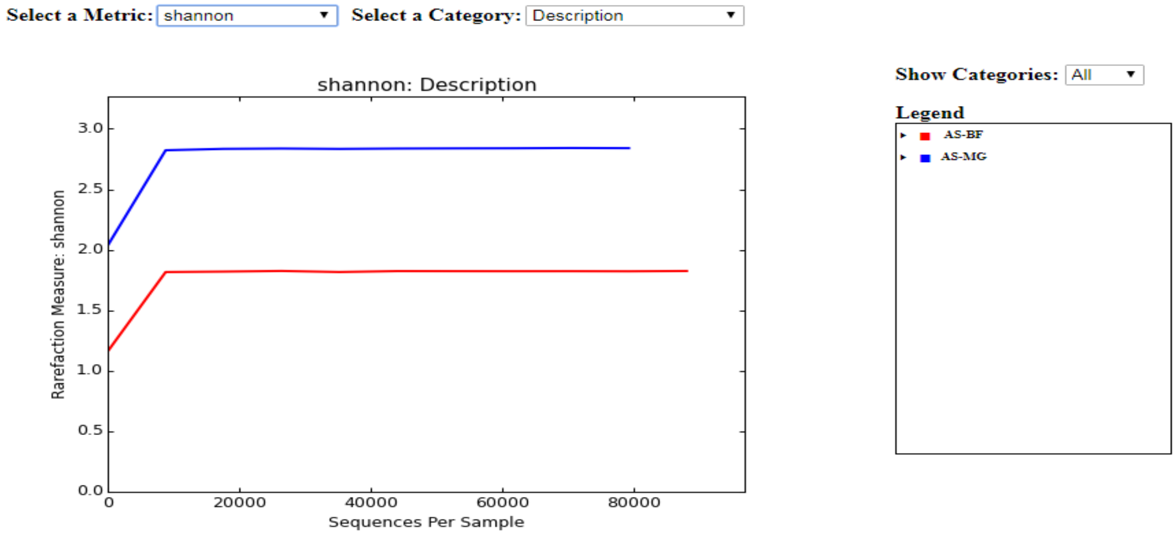


**Fig. S5:** Graphical representation of the diversity indices (a) Chao1 (b) Shannon alpha-diversity rarefaction curves of the sugar fed and blood fed midgut microbiomes of the *Anopheles stephensi* showing full extent of phylotype richness and quantifiable diversity estimation.

**FIG. S6 a & b**

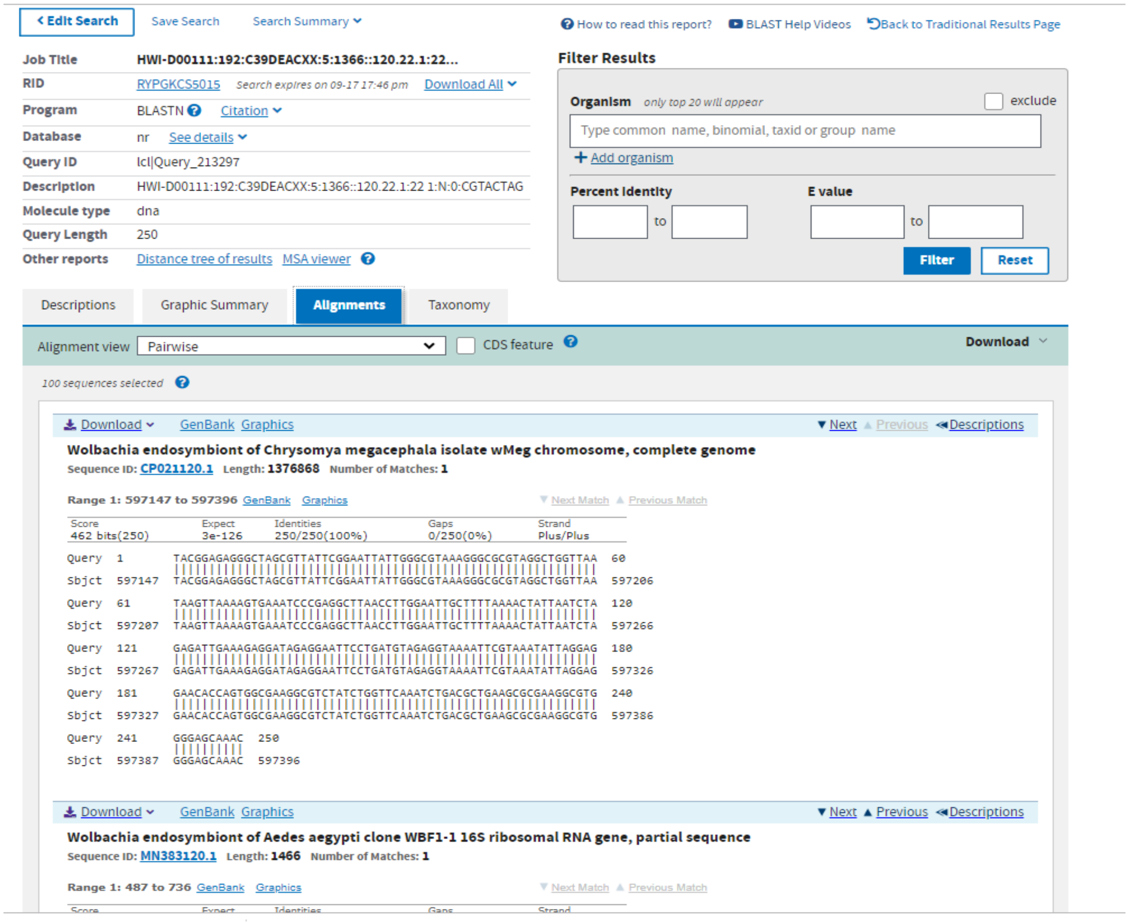


**Fig S6a:** A 250 bp long Metagenomic reads NCBI/BLASTn analysis against NR database identifies Wolbachia endosymbiont sequnce in the mosquito *Anopheles stephensi* gut.

1. AS_MG_SF

CDS_1174_Transcript_250

hypothetical protein[Wolbachia endosymbiont of Drosophila ananassae]

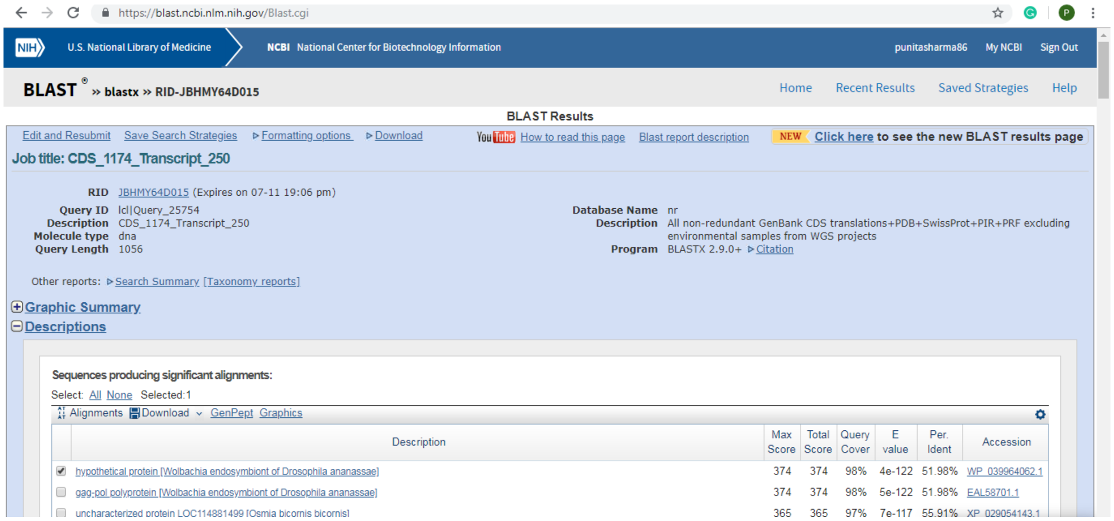


2. CDS_7405_Trasncript_21832

CDS_7405_Trasncript_21832
hypothetical protein [Wolbachia pipientis]

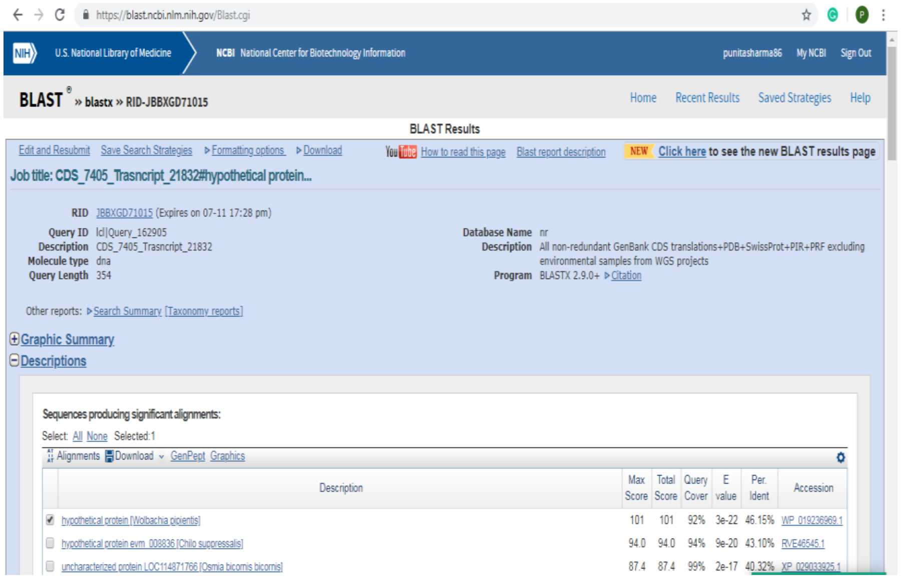


**Fig. S6b:** RNAseq based BLASTx analysis of putative trancripts predicts  Wolbachia endosymbiont homolog proteins in the mosquito *Anopheles stephensi* gut.

Fig S7

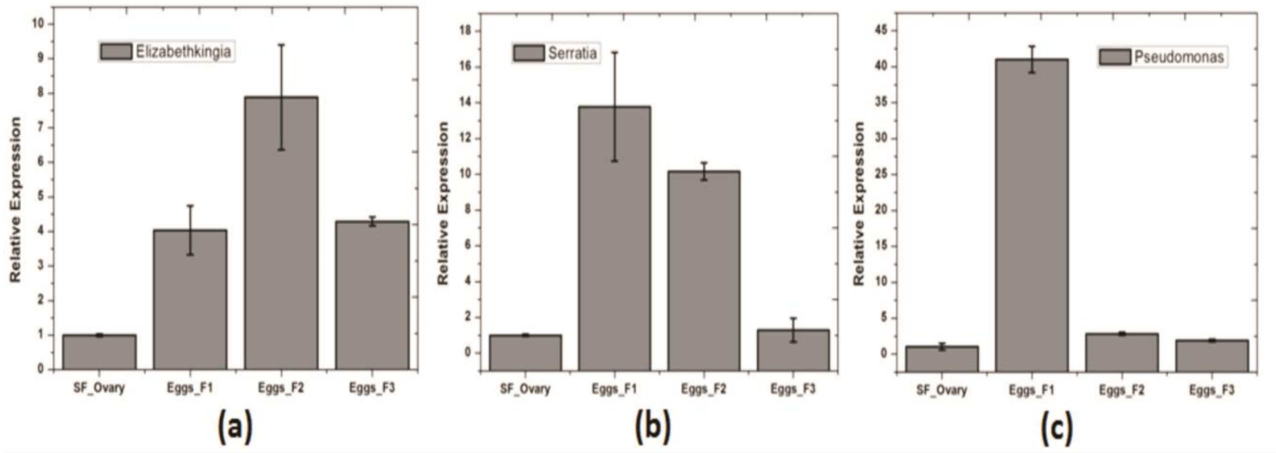


**Fig. S7**: Relative quantity of different bacteria in the *Anopheles stephensi* ovary and the eggs of subsequent generations**:** In this figure the relative abundances of the selected bacteria viz. *Elizabethkingia*, *Serriatia* and *Pseudomonas* in the ovary of the parent generation and then first batch of eggs of the subsequent generations of the mosquito (F1, F2, F3) were relatively quantified.
