## SUPPLEMENTALE TABLE FILE_S1_S2_S4_S6_S7 for "Altered gut microbiota and immunity defines *Plasmodium vivax* survival in *Anopheles stephensi*"

**ST 1: Primer sequences used for the amplification of the V3-V4 hypervariable region of 16S rDNA gene of Eubacteria and Archaea for the 16S metagenomic library preparation**

| Sr.No. | Oligo Name | Oligo Sequence ( 5' to 3') | Length of primer | Product size (Approx.) |
|---|---|---|---|---|
| 1 | V3-Forward | CCTACGGGNGGCWGCAG | 17 | ~ 460 bps |
| | V4-Reverse | GACTACHVGGGTATCTAATCC | 21 | |

**ST-2 List of the primers used in the Real time and RT- PCR**

| Sr. No. | Primer name | Forward Primer sequence (5' to 3') | Reverse Primer sequence (5' to 3') |
|---|---|---|---|
| 1 | Elizabethkingia | CGAGCGGTAGAGATTCTTCGG | ACACGTAAGTAGGTTTATCCCCAGA |
| 2 | Pseudomonas | GACGGGTGAGTAATGCCTA | CACTGGTGTTCCTTCCTATA |
| 3 | Serratia | CTGTCGTCAGCTCGTGTTGT | TTCATGGAGTCGAGTTGCAG |
| 4 | 16s | TTGGAGAGTTTGATCCTGGCTC | ACGTCATCCCCACCTTCCTC |
| 5 | C1 | AACCAACCGAACCGTATCAA | TTTCTCAGCTGCCTTGAACA |
| 6 | C2 | AAGCTGCTCTTTCTCGTTGC | GTGAGGTACGCCCTATCCAA |
| 7 | D1 | GATGAACTGCCCGAAGAGAC | TTGCTGGCTGTTGCAGTATC |
| 8 | Gambicin | ACTGTGGCTACGGGTACGTC | GCTTGTTCTTCCGGTGTGAT |
| 9 | Lem A | CGTTTACCAGAAACGTGCAA | TGCTGGTCTGCCTTTAGGTT |
| 10 | FECR | TTTCTTCCGGCTTTTGGATA | AAATATTGCAGTCCCGCTTG |
| 11 | ABC transporter | TTGTATCGATCATGGGGTCA | TCTTTTCGGGAAACATTCTGA |
| 12 | Sus C | TAGATGCGAACGGACTTCCT | CGGTTCCATCAGCAACTACA |
| 13 | Ton B | CATTGGGAAAGTAGGCGTGT | GACTGGATCCTGGCTTACCA |

**Supplemental Table ST4: Blood fed mosquitoes gut RNAseq database assembly and NR homology search analysis statistics**

| Description | ASMGC |
|---|---|
| Total No. of bases | 35,072,112 |
| No. of Contigs | 74,548 |
| Total no. of transcripts | 11159 |
| Average Transcript Size | 470 |
| N50 | 539 |
| Max Transcript Size | 5,536 |
| Total CDS analyzed (Post duplicate removal) | 7,381 |
| Best match to NR database | 5042 |
| Insect homolog sequences | 4,562 |
| Unknown/Hypt | 2340 |
| Non insect microbial | 479 |

**ST-6: Table showing the putative function of bacterial (EK) genes present in the *Anopheles stephensi* transcriptome and showed enriched expression in response to blood feeding.**

| Gene | Putative function |
|---|---|
| LemA | Quoram sensing, biofilm production,interspecies communication and siderophore production |
| Ton B,<br>FecR,<br>ABC transporter | Siderophore uptake and iron metabolism |
| SusC/RagA | import large degradation products of proteins (e.g. RagA) or carbohydrates (e.g. SusC) as nutrients |

**ST-7 Details of the *Wolbachia* transcript coding protein retrieved from the different transcripptomes of  *Anopheles stephensi*  midgut.**

| Sr. no. | Transcript label | Origin | Size (bp) | Coding nature | Percentage identity |
|---|---|---|---|---|---|
| 1. | CDS_1174_Transcript_250 | AS_MG_SF | 1056 | hypothetical protein[Wolbachia endosymbiont of Drosophila ananassae] | 51.98% |
| 2 | CDS_3077_Trasncript_20360 | AS_MG_BF | 546 | hypothetical protein [Wolbachia endosymbiont of Drosophila ananassae] | 43.81% |
| 3 | CDS_7405_Trasncript_21832 | AS_MG_PV_6-8D | 354 | hypothetical protein [Wolbachia pipientis] | 46.15% |
| 4 | CDS_7433_Trasncript_21981 | AS_MG_PV_6-8D | 396 | hypothetical protein [Wolbachia endosymbiont of Drosophila simulans] | 31.88% |
| 5 | CDS_11533_Trasncript_48370 | AS_MG_PV_6-8D | 342 | hypothetical protein [Wolbachia endosymbiont of Drosophila simulans] | 43.01% |
| 6 | CDS_698_Trasncript_4595 | AS_MG_PV_10D | 1092 | hypothetical protein [Wolbachia endosymbiont of Drosophila simulans] | 32.45% |
| 7 | CDS_5445_Trasncript_19039 | AS_MG_PV_10D | 426 | hypothetical protein [Wolbachia pipientis] | 33.33% |
